# Structure-Led Exploration of the Metagenome Yields Novel RNA-Guided Nucleases with Broad PAM Diversity

**DOI:** 10.64898/2026.03.27.714800

**Authors:** Emmanuel LC de los Santos, Lila Rieber, Meng Wang, Seana Catherman, Stephen Hatfield, Tyson Bowen

## Abstract

Compact RNA-guided nucleases with favorable targeting properties are challenging to discover due to their low natural abundance. Here, we develop a structure-led search strategy -leveraging predicted protein folds and sequence-independent similarity metrics -to systematically identify extremely low-homology compact RNA-guided nucleases across vast metagenomic datasets with high computational efficiency. Homology clustering resolved these proteins into distinct groups, for which we performed comprehensive PAM profiling and evaluated editing efficiency in eukaryotic cells. This structure-guided discovery revealed a previously undiscovered landscape of compact nuclease subtypes that exhibit extensive protospacer-adjacent motif (PAM) diversity, expanding the targeting potential of compact editors.

Comparative analysis across the novel RNA-guided nuclease families demonstrates that compact systems are not intrinsically limited to highly constrained PAMs but instead have a broad and previously unknown breadth of genome targeting capabilities, comparable to that of Cas9 and far exceeding common transposon- derived systems. Additionally, this search revealed that a compact transposon-associated motif (TAM) is a prerequisite for the emergence of a CRISPR-Cas system from ancestral transposons, before protein domain expansions increase the target length and specificity constraints. These results enrich the catalog of RNA-guided nuclease architectures and contribute validated compact genome editing tools with broad and diverse PAM recognition, which may have therapeutic applications.

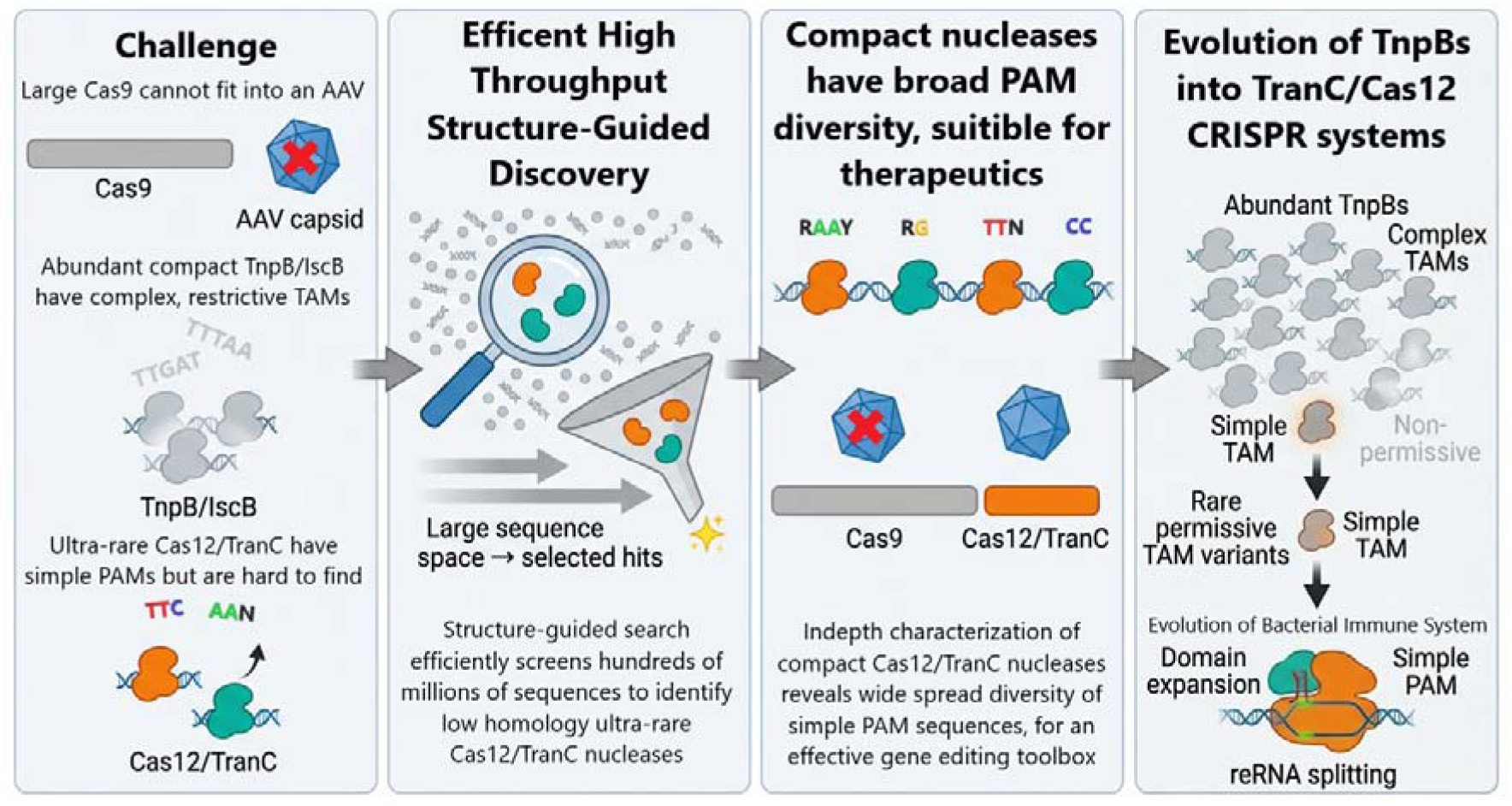

## Background

RNA-guided nucleases have transformed genome engineering, enabling programmable manipulation of DNA for research and therapeutic applications^1,2^. However, widely used systems such as Cas9 remain constrained by their large size, which limits efficient delivery in vivo, particularly for viral vectors^3,4^. In contrast, compact RNA-guided nucleases derived from TnpB and IscB are highly abundant in microbial genomes but typically require long and restrictive target-adjacent motifs (TAMs), limiting their practical targeting scope^5,6^. Compact Cas12-related systems offer a potential solution, combining smaller size with simpler PAM requirements; however, these systems appear to be comparatively rare and thus difficult to identify using conventional sequence homology–based approaches^4,7–11^. Recent work has identified TranC nucleases as evolutionary intermediates between TnpBs and Cas12 systems, providing insight into the origins of RNA-guided nucleases^12^. Notably, the systems characterized in this study were identified independently through structure-guided searches prior to the availability of that classification. Despite these advances, the diversity, targeting capacity, and genome editing potential of compact Cas12/TranC nucleases remain largely unexplored.

Here, we address this gap by combining structure-guided discovery with systematic functional characterization of compact RNA-guided nucleases (RGNs) across diverse subtypes. By enabling scalable exploration of hundreds of millions of candidate proteins beyond the limits of sequence homology, our approach identified diverse compact systems with minimal sequence similarity to previously characterized nucleases and independently discovered the recently reported TranC evolutionary intermediates between TnpB and CRISPR-Cas systems. We show that compact nucleases exhibit unexpectedly broad protospacer adjacent motif (PAM) diversity and targeting capacity, in contrast to the restrictive TAM requirements of TnpB- and IscB-derived systems. Furthermore, we demonstrate efficient genome editing in mammalian cells, including primary human T cells, and establish their specificity relative to existing compact nucleases. These results provide evidence that a compact TAM/PAM acts as an evolutionary filter along the path from TnpB to TranC to Cas12 RGNs. Our findings demonstrate compact Cas12/TranC systems are a versatile and practical class of genome editors with expanded targeting potential for the basis of an effective therapeutic tool box^13^.

CRISPR-Cas systems are adaptive immune systems in bacteria, archaea, and phage genomes. Their ability to cleave double-stranded DNA (dsDNA) using a reprogrammable guide RNA has been developed into a variety of genome editing tools in the past decades^1,2^. CRISPR-Cas systems are divided into two distinct classes based on the organization of their effector modules. Class 1 systems employ a multi-subunit Cas protein complex, while Class 2 systems utilize single effector proteins, including the well-characterized Cas9 (Type II) and Cas12 (Type V)^14–18^.

Type II and most Type V RGNs precisely target dsDNA for cleavage by binding a reprogrammable target RNA molecule, consisting of a CRISPR RNA (crRNA) and often a trans activating CRISPR RNA (tracrRNA), to a complementary DNA sequence adjacent to a PAM. While the target sequence may be changed to almost any sequence, the PAM recognition is an intrinsic attribute of the protein sequence and limits the targeting ability of the Cas enzyme to regions containing the PAM^5^.

This restriction has been the primary cause for the ongoing search for novel CRISPR systems better suited to therapeutic editing^13^. As one example, our team’s recent research into generating allogenic (“off-the-shelf") CAR-T cells resistant to both biochemical (adenosine) and immunological (PD-L1 and TGF-β) inhibitory signaling for improved activity in solid tumors required the use of a Cas9 base editor and an additional RGN, due to the limits on targeting imposed by PAM and precision editing requirements^19–26^.

Type II systems, which have 3’ PAMs, are abundant in microbes, accounting for 24% of bacterial CRISPR systems, and have diverse PAM sequences^4^. In contrast, Type V systems, which have 5’ PAMs, comprise only 6% of CRISPR systems, and 81% are the Type Va (Cas12a) subtype^4^. Cas12a proteins recognize a much smaller range of PAM sequences, typically 5’ TTN^4^. Other, rarer Type V subtypes, such as Cas12f and Cas12n (originally called c2c9), exhibit potentially greater PAM diversity, but their investigation has been limited by their relative paucity in microbial genomes^7,27,28^.

In contrast, the compact ancestral proteins of the effector proteins for Class II CRISPR systems – IscB for Type II systems and TnpB for Type V systems – have varying degrees of recognition sequence diversity^5,6,29–31^. As each one of these ancestral proteins represents a potential novel origin for the evolution of CRISPR systems, it is possible that further compact CRISPR systems with diverse PAMs remain to be discovered. Indeed, recent work has identified reprogrammable transposon-CRISPR intermediates (TranC) nucleases as early steps in the transition from TnpB transposons to Cas12 CRISPR-based immunity nucleases, which are likely to have arisen from multiple independent TnpB origins^12^.

Discovering novel reprogrammable effector proteins requires the identification of nucleases or CRISPR-associated (Cas) proteins near a CRISPR repeat array. The first Cas proteins were identified by finding homology across species among the open reading frames (ORFs) adjacent to CRISPR arrays^32^. Many computational search strategies have since been developed to use sequence homology from the multitude of known Cas protein sequences to rapidly identify and characterize more potential candidates. These approaches tend to search for multiple Cas proteins in an operon, a computationally expensive strategy, especially given the rapid growth of sequence databases^33,34^. However, smaller CRISPR-Cas operons that consist only of a single effector protein and a CRISPR array could be missed by this method, especially for rare subtypes^8^. In addition, these searches assume that novel proteins have at least moderate homology to known proteins, limiting the ability to discover extremely divergent subtypes, or subtypes that have evolved from a different ancestral TnpB. New subtypes represent an ever smaller fraction of the total CRISPR landscape, making their identification more challenging, even with the proliferation of sequence databases^18^. While more computationally efficient approaches, such as efficient retrieval-augmented search tool (ERAST) and fast locality-sensitive hashing–based clustering (FLSHclust), have been developed and, in the case of FLSHclust, used to discover previously unreported CRISPR-linked systems, they still rely on sequence homology to identify novel systems^10,35^. Novel Cas enzymes with low homology to known enzymes can be identified by sequencing large microbial and metagenomic environments to increase confidence in their co-occurrence with CRISPR repeats, but this requires extensive sequencing, constructing search profiles from dozens to hundreds of representative sequences, and a subsequent analysis effort that can still miss subtypes below a certain abundance or similarity threshold^11,12,36,37^. These limitations call for different approaches to identifying novel systems, especially for searching collections where additional sampling is impractical or impossible, such as public databases.

Unlike Cas9, for which all known variants appear to have evolved from a single IscB ancestor, convergent evolution of multiple different TranC and Type V subtypes has occurred from independent ancestral TnpB sequences, leading to their impressive sequence and functional diversity, but also creating challenges for sequence-based searches^12,38^. Rather than relying on increasingly remote sequence similarity, we sought to search for new effector proteins and subtypes using structural similarity, which we hypothesized would lead to functional similarity, even with extremely low sequence homology^39,40^. Structural searches have been used to discover novel orthologs of enzymes with little previously known sequence diversity, including the ancestral clade of CRISPR-Cas13 ribonucleases, and functions have been predicted from diverse primary sequences by clustering predicted structures^40–42^.

Recent advances in machine learning have revolutionized protein structure prediction, with tools such as AlphaFold2 allowing the rapid prediction of three-dimensional structure of proteins^43^. The release of ESMFold, which uses embeddings from a protein language model to predict structure, rather than computationally expensive multiple sequence alignments, allowed for structural predictions at an even greater scale^44^. This was highlighted by the release of over 600 million ESMFold predicted metagenomic protein structures, which we searched against. We supplemented these structural databases with an internal collection of ESMFold predicted structures from ORFs near CRISPR arrays in the human gut microbiome, which has been shown to be enriched for microbes with CRISPR systems, as well as in viral databases, which have been shown to possess unusual CRISPR subtypes, and a collection of microbes from diverse microbiomes^4,45–49^. We then compared the characteristic RuvC-like nuclease (RuvC) domain fold of a few Class 2 CRISPR effector proteins to these databases using Foldseek, a method developed to rapidly search structural databases^50^. These tools enabled the search of large databases in structural space in a computationally tractable manner, by flattening the three-dimensional structural components in a meaningful way to enable highly efficient one-dimensional search algorithms, in contrast to the CRISPR-Cas13 search which relied on older, much more computationally intensive tools for structural comparison^40^. Our approach led us to discover new examples of CRISPR systems of both known rare subtypes and novel subtypes using only four search queries, without creating novel motif searches, profile searches, or sequence searches from dozens or hundreds of representative systems, as is typically performed11,12,37,38,51.

Correct identification of the RNA components of the putative CRISPR systems is required for validation and functional characterization. Again using the hypothesis that structural conservation leads to shared activity, we predicted tracrRNA sequences for compact Type V effector proteins, including Cas12f, Cas12n, and two TranC clades^7,27^. The ancestral Cas12 protein, TnpB, can self-process its mRNA to generate guide ωRNA, located in the ORF^29^. This also occurs to the tracrRNA of Cas12n systems, which are thought to be early evolutionary intermediates of Type V systems^27^. Both guide RNA sequences have predicted minimum free energy (MFE) structures similar to the tracrRNA-crRNA also identified in Cas12f systems. This conserved MFE structure, combined with incorporating the antirepeat-repeat hybridization of Cas12 crRNA sequences, can be used to predict the tracrRNA for many compact Type V subtypes, analogous to what has been done for Cas9^28,52,53^.

Using this approach to predict tracrRNA sequences for our discovered CRISPR systems, we functionally characterized dozens of these compact systems, in both bacteria and eukaryotes, and compared the PAM diversity of large numbers of these effector proteins. We demonstrate that compact Cas12 and TranC systems have a large degree of PAM sequence diversity, unlike Cas12a, and, unlike the highly abundant TnpB and IscB ancestral sequences, their PAMs are short and enable target flexibility. We further show that these same compact systems generally have a higher specificity than TnpB and demonstrate their ability to create T cells that are resistant to biochemical and immunological inhibitory signaling. In summary, our approach rapidly and easily identified novel highly active compact genome editors with low sequence homology to known Cas12 effector proteins and with diverse PAM requirements, illustrating the convergent evolution of different TnpB ancestral proteins into diverse Type V CRISPR systems and their applicability for human genome engineering.

## Results

### Computational Search Strategy

To identify novel RGN sequences, we focused on identifying proteins near a CRISPR array that contain the ancestral RuvC domain, as this fold was thought to be present in the last common ancestor of both IscB and TnpB^54^. As a basis for the RuvC search, we extracted the coordinates of the Pfam RuvC fold and the RuvC folds of four RGNs with crystal structures (SpCas9 (5F9R), FnCas12 (5NFV), Cas12f1 (7C7L), and CasLambda (8DC2)), representing a wide sequence and structural diversity (Supplemental Figure 1). ^55–59^ The RuvC domain of CasLambda in particular is so divergent from other examples that it was not readily identifiable from the primary sequence, although its biochemical and structural similarity led it to be classified as a Cas12 enzyme^59^. When comparing the primary sequence or structural alignments, the structural similarity of the RuvC domain exhibited higher similarity among the four RGNs than the sequence or the entire structures. As such, the smaller size of the extracted RuvC domain was ideal to use as an efficient search parameter to identify systems with extremely low sequence homology. The extracted RuvC domains were used in a Foldseek search against a compiled database of ESMFold predicted protein structures. To determine suitable cutoffs for the Foldseek e-values, we first conducted a search on a set of ESMFold predictions of known RGNs, selecting an e-value cutoff such that each RGN sequence was a hit to least one of the four RGN RuvC folds used.

Several datasets were used for the Foldseek query, including the Foldcomp-compressed full ESMAtlas database and ESMFold predicted structures for coding sequences within 10 kilobases of a CRISPR array from gut and environmental bacterial, archaeal, and viral metagenomic sequences (Figure 1a). Foldseek hits of the ESMAtlas were traced back to their source genomic DNA to identify hits that were proximal (<10kb) to a CRISPR array, though this was often not possible due to limitations in the metadata. Filters were applied to reduce the number of candidates, such as removing sequences that were too large, truncated, or that did not have the catalytic DED residues of RuvC. The resulting sequences were highly divergent from the full-length sequence of the initial four search systems and represented a wide diversity of sequences.

**Figure 1.**
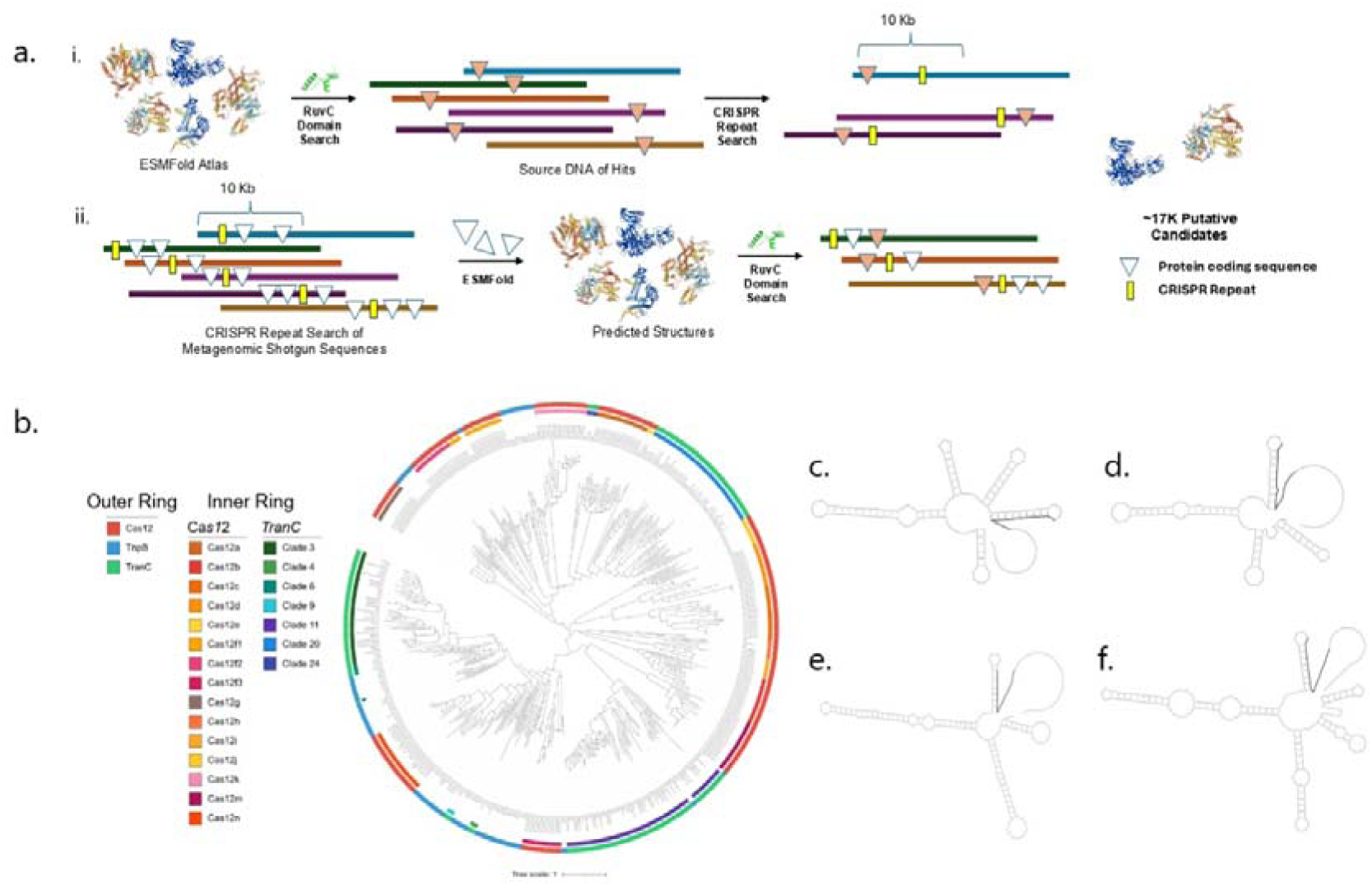
Structure-driven search strategy for novel nucleases and associated RNA elements. a.i. “*Structure-first search*” Foldseek was used to identify the presence of the RuvC domain on predicted structures of metagenomic sequences from the ESMAtlas. Coding sequences that contained a CRISPR repeat in proximity of a RuvC hit were carried forward as candidate nucleases. a.ii. “*Sequence-first search”* CRISPR repeats were identified in DNA sequences from metagenomic shotgun sequencing databases. Coding sequences close to CRISPR repeats whose predicted structures contained RuvC domains were considered as candidate nucleases. b. Phylogenetic tree of rare and novel nucleases identified from structure driven search compared to known examples of RuvC containing nucleases. c-f. MFE diagram or representative gRNA of Cas12n(c), Cas12f(d), TranC clade 3(e), TranC clade 11(f). Highlighted region is the repeat sequence of the crRNA.

To select putative systems for experimental characterization, we compared the structure predictions to known RGNs. This was done by doing an all-versus-all alignment using Foldseek and using a distance metric based on pairwise template modeling scores, a measure of the structural alignment between two protein structures. This enabled us to assign putative classes to the CRISPR systems we discovered. We excluded systems that were identified as non-cutting subtypes (e.g. Cas12k or Cas12m), IscB-like, or similar to known RGNs (>80% sequence identity), and focused on compact CRISPR systems that were new examples of rare subtypes or systems from subtypes undiscovered at the time of our search^60,61^. We removed systems for which proximity to the CRISPR array appeared coincidental rather than functional by excluding those with highly inconsistent distance or positioning relative to the array. Finally, we augmented this list by doing a sequence-based similarity search to identify more putative CRISPR systems of our selected types.

Overall, from rare subtypes we selected one Cas12b, one Cas12e, three Cas12h, 25 Cas12f, 34 Cas12n, and 54 sequences from multiple novel subtypes for characterization^7,9,27,62,63^. After our search, the characterization of TranC clades was published, and our novel subtypes correspond to the previously characterized TranC clades 3 and 11, as well as the putative clades 4, 6, 9, 20, and 24^12^. The phylogenetic comparison of the various subtypes of Cas12, TranC, and TnpB does not show consistent out groupings of any one subtype but instead shows close evolutionary relationships between the different Cas12 and TranC subtypes, interspersed with various TnpB family members, reflecting the multiple independent evolutionary events (Figure 1b). We identified systems from bacteria, archaea, and viral genomes across multiple different subtypes (Supplementary Table 1).

### Identification and Comparison of tracrRNA structure

All the known subtypes we identified, except for Cas12h, require a tracrRNA^60^. The tracrRNA for Cas12b and Cas12e were readily determined by identifying an antirepeat and an upstream sequence that conformed to previous reports of tracrRNA structures for those subtypes^64,65^. It has been previously reported that the Cas12n subtype possesses an unstructured and dispensable C-terminal tail that contains the tracrRNA^27^. We aligned the Cas12n sequences and compared the predicted local distance difference test (pLDDT) values, a measure of structure confidence, across the alignment to identify the putative tracrRNA for Cas12n (Supplemental Figure 2)^66^.

We observed a consistent drop in structure confidence after the final catalytic RuvC residue across the Cas12n subtype, consistent with the reported unstructured tails. We then identified the nearest antirepeat to the end of the low-pLDDT region, which gave a putative tracrRNA of between 100-200 bp without incorporating any of the 10 amino acids after the final catalytic RuvC aspartic acid. When joined to the appropriate putative crRNA sequence from the CRISPR array, we observed a consistent structure of the gRNA of approximately three to four stem loops and the final antirepeat-repeat stem, followed by a short stretch of unpaired “leader” bases before the reprogrammable target sequence (Figure 1c). This structure showed remarkable similarity to published gRNAs of Cas12f enzymes^28,67,68^. As Cas12f does not have an unstructured tail, we searched for an antirepeat sequence between the effector protein sequence and the CRSIPR array between 100-200 bases that would adopt a consistent structure (Figure 1d)^27^. None of the novel subtypes showed an unstructured tail like Cas12n (data not shown), so we searched in a similar manner to Cas12f systems for a tracrRNA sequence and were able to identify a consistent structure in two of the TranC clades but not in the others (Figure 1e-f).

### Functional Characterization

#### PAM Activity and Diversity

The rare and novel subtypes were tested against an exhaustive 8mer PAM library positioned 5’ to the guide sequence. We identified the PAM for the Cas12b, Cas12e, two Cas12h (67%), 21 Cas12f (84%), and 18 Cas12n (53%), 15 TranC clade 11 (71%), 9 of TranC clade 3 (90%), 4 of the novel TranC clade 20 (31%) and the novel TranC clade 9 representative, but not for clades 4, 6, or 24. (Active sequences in Supplemental Table 1, PAM sequences in Supplementary Figure 3). We saw a higher success rate in those subtypes for which we codon-optimized for expression in *E. coli* and expressed the gRNA from a separate T7 promoter compared to expressing from the CRISPR operon without any optimization. This success rate was comparable with our previous work on identifying diverse Cas9 (68%) and Cas12a (63%) sequences, demonstrating the robust applicability of our method to identify novel active systems^4^. Surprisingly, the first Cas12h system we tested did not function when codon-optimized with separate expression of the expected crRNA but did function as the native CRISPR operon, despite the reported dispensability of everything except the gene sequence and the CRISPR array^63^. The remaining two Cas12h systems were only tested as native CRISPR operons.

We next calculated a targetability score for each system and relevant reference systems, to compare the ability of each system to target across a hypothetical genome^4,5,12,29,30,69^. The score represents the number of times that PAM would be expected to be found in a sequence of 1000 bases with an even distribution of all four nucleotides on one strand (Figure 2a). We then calculated median Euclidean distance (MED) of recognition sequences for each subtype to measure the diversity of sequences that they can target (Table 1). The MED score represents the pairwise distances among recognition sequences, with higher scores indicating more diversity. We compared the newly identified rare and novel compact PAMs alongside validated Cas9 and Cas12a PAMs to understand the full range of potential target sequences of each subtype (Figure 2b)^4,12,69^. Cas12a shows much less PAM sequence variation than Cas9 but a similar range of sequence targetability per PAM. In contrast, the rare and novel compact systems demonstrate a range of PAM sequence diversity almost comparable to that seen in Cas9, but with an equal or greater sequence targetability than seen in Cas9 or Cas12a. We then compared the PAM diversity to the reported transposon-associated motif (TAM) sequences from IscB and TnpB systems (Figure 2c)^5,30^. There is a greater diversity in IscB TAM sequences than Cas9 PAMs but the TAMs exhibit a much smaller targetability than Cas9 PAMs. In contrast, there is much greater diversity in TnpB TAMs than in Cas12a PAMs but still a drastically smaller targetability with TnpB TAMs. The diversity of the TnpB TAMs is similar to what is observed with the rare and novel compact system PAMs, however. However, the PAMs of the rare and novel compact systems exhibit a much higher targetability than the TnpB TAMs and there is a range of recognition sequence diversity within each subtype.

**Figure 2.**
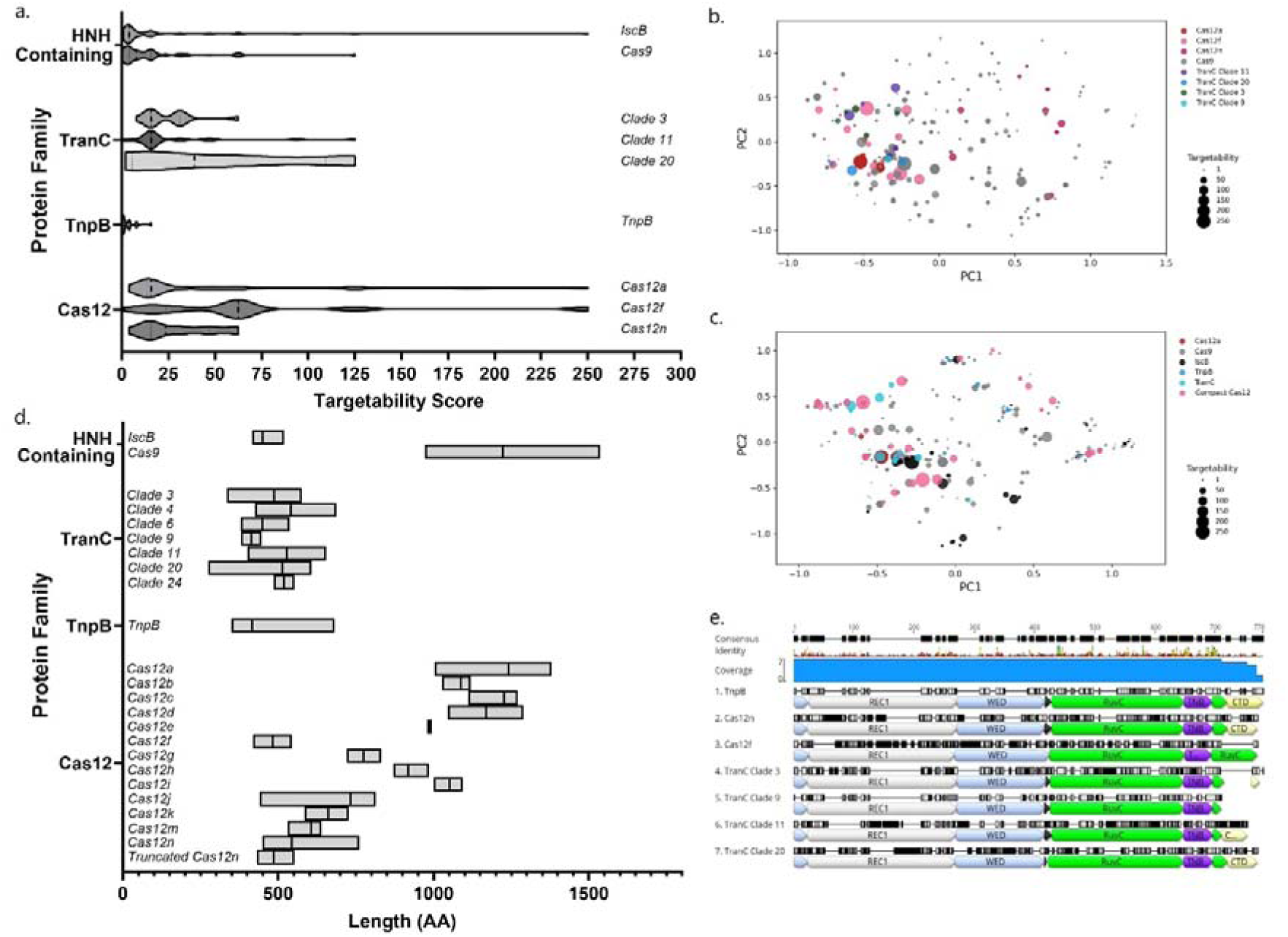
Recognitions Sequence and Protein Characteristics of Novel and Compact Systems. a. Distribution of targetability scores of newly discovered rare and novel RGNs compared to Cas9, IscB, Cas12a, and TnpB. Higher values indicated a more frequently occurring recognition sequence. Median is designated with thicker dashed line, quartiles with thin dashed lines. b. Principal component analysis of PAM depletion scores of newly discovered rare and novel RGNs compared to Cas9 and Cas12a. Increasing marker size corresponds to a greater targetability score. c. Principal component analysis of reported PAM and TAM sequences of RGNs and ancestral transposons. Increasing marker size corresponds to a greater targetability score. d. Distribution of sizes of RGNs. E. Domain organization of representative active effector proteins relative to ancestral TnpB protein.

**Table 1.**
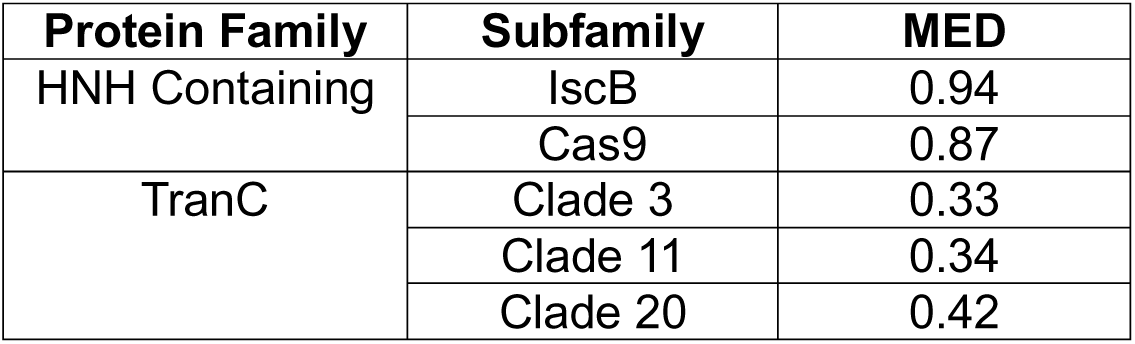
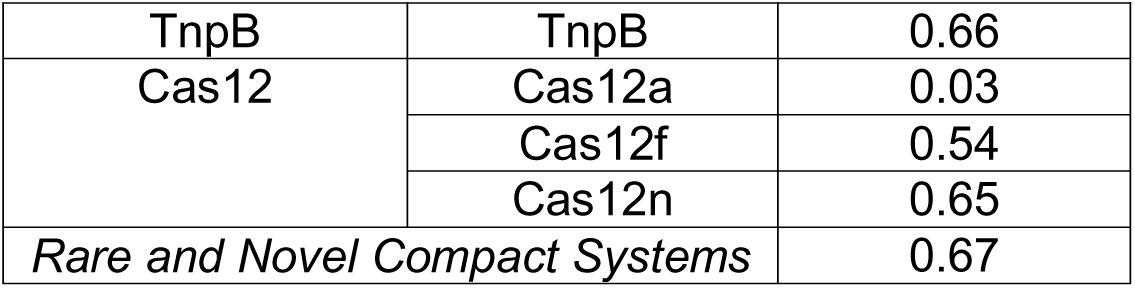
Median Euclidean distance of PAMs and TAMs of RGN Families.

| Protein Family | Subfamily | MED |
| --- | --- | --- |
| HNH Containing | *IscB* | 0.94 |
|  | Cas9 | 0.87 |
| TranC | Clade 3 | 0.33 |
|  | Clade 11 | 0.34 |
|  | Clade 20 | 0.42 |
| TnpB | TnpB | 0.66 |
| Cas12 | Cas12a | 0.03 |
|  | Cas12f | 0.54 |
|  | Cas12n | 0.65 |
| *Rare and Novel Compact Systems* |  | 0.67 |

#### Subtype Comparison

The newly identified representatives of known Cas12 effector proteins and TranC clades conformed to the expected size range of their subtype (Figure 2d). Removing the unstrucutred tail from Cas12n changed the average size from 544 amino acids to 485 amino acids, representing a reduction of approximately 10% of the protein. This reduction brings them close to the average length of Cas12f proteins (481 amino acids) but still noticably larger than the average length of the ancestral TnpB (415 amino acids). TranC Clade 20 has an average length of 513 amino acids and Clade 9 contains the shortest nucleases, with an average length of only 414 amino acids, almost equal to the ancestral TnpB length. This clade appears to be the smallest CRISPR-associated nuclease with demonstrated activity, comparable to the smallest active TnpB proteins. Cas12f and Cas12n have expanded REC and WED domains compared to TnpB^70^. When compared, by either sequence or predicted structure, to TnpB (PDB ID: 8H1J), Cas12f (PDB ID: 7C7L), or Cas12n (PBD ID: 9J09), our active

TranC9 protein was identified in a caudoviricetes genome and does not appear to have the CTD from ISDra2TnpB, nor the expanded REC and WED domains of Cas12f and Cas12n^57,70,71^. The predicted structure shares high similarity to ISDra2 TnpB, and will be refered to as CauTranC9 (Supplemental Figure 4). As previously reported, TranC Clade 3 is similar to ISDra2 TnpB, posessing just an enlarged bridge helix within the RuvC domain^12^. In contrast, TranC Clade 11 and Clade20 have expanded REC and WED domains, in addition to an elongated CTD like that seen in TnpB (Figure 2e). This CTD is not comparable to the unstructured CTD of Cas12n, as the tracrRNA is not found in this region, and the pLDDT values of the CTD are high, indicating a consistent predicted structure (data not shown).

The 54 sequences from the novel subtypes were searched for nearby Cas accessory proteins with CRISPRCasTyper^33^. The only nearby accessory proteins identified belonged to other known subtypes of CRISPR systems and were not associated with the novel effector protein or CRISPR array. The CRISPR arrays were primarily predicted to belong to an unknown subtype, and the repeats were 32 or 36 bp long for TranC clade 3 (median 32 bp), 38 bp for clade 9, 32-36 bp for clade 11 (median 33 bp), and 36 or 38 bp long for clade 20 (median 36 bp). For the arrays not near the end of a contig, there were 3-20 repeats for clade 3 (median 4.5), 5 repeats for TranC clade 9, 3-17 for TranC clade 11 (median 5), and 3-12 for Clade 20 (median 5). The native CRISPR array of the ultra-compact CauTranC9 targets phage sequences (Supplementary Table 5) indicating it is functioning as an antiphage defense as expected^48,72^.

#### Programmable Eukaryotic DNA Editing

The systems with an identifiable tracrRNA and a confirmed PAM sequence were tested in HEK293T cells for reprogrammable eukaryotic genome editing. Consistent with previous results, many of the systems demonstrated low, but detectable, editing in eukaryotic cells (Supplemental Figure 5)^3,4,60,73^. Six compact systems with the highest initial activity were selected for further evaluation: two Cas12f systems (from *Alistripes* genus and from the Desulfovibrionaceae family) - named AlCas12f and DeCas12f, two truncated Cas12n systems from the Rothia genus (*Rothia dentocariosa* and Rothia sp. HMSC069C03) - named Rd2Cas12n and RoCas12n, a member of the TranC clade 11 from *Ruminococcaceae bacterium* (RuTranC11), and a member of the TranC clade 3 from *Oscillospiraceae HGM13006 sp905202605* (OsTranC3). These compact rare or novel systems were further improved when possible by truncating the gRNA, consistent with previous reports (Supplementary Figure 6)^27,67,68^. The optimal gRNA structure was then used to evaluate the ability of these systems to perform genome editing in eukaryotic cells compared to AsCas12a across 12 targets in triplicate (Figure 3a)^17^.

**Figure 3.**
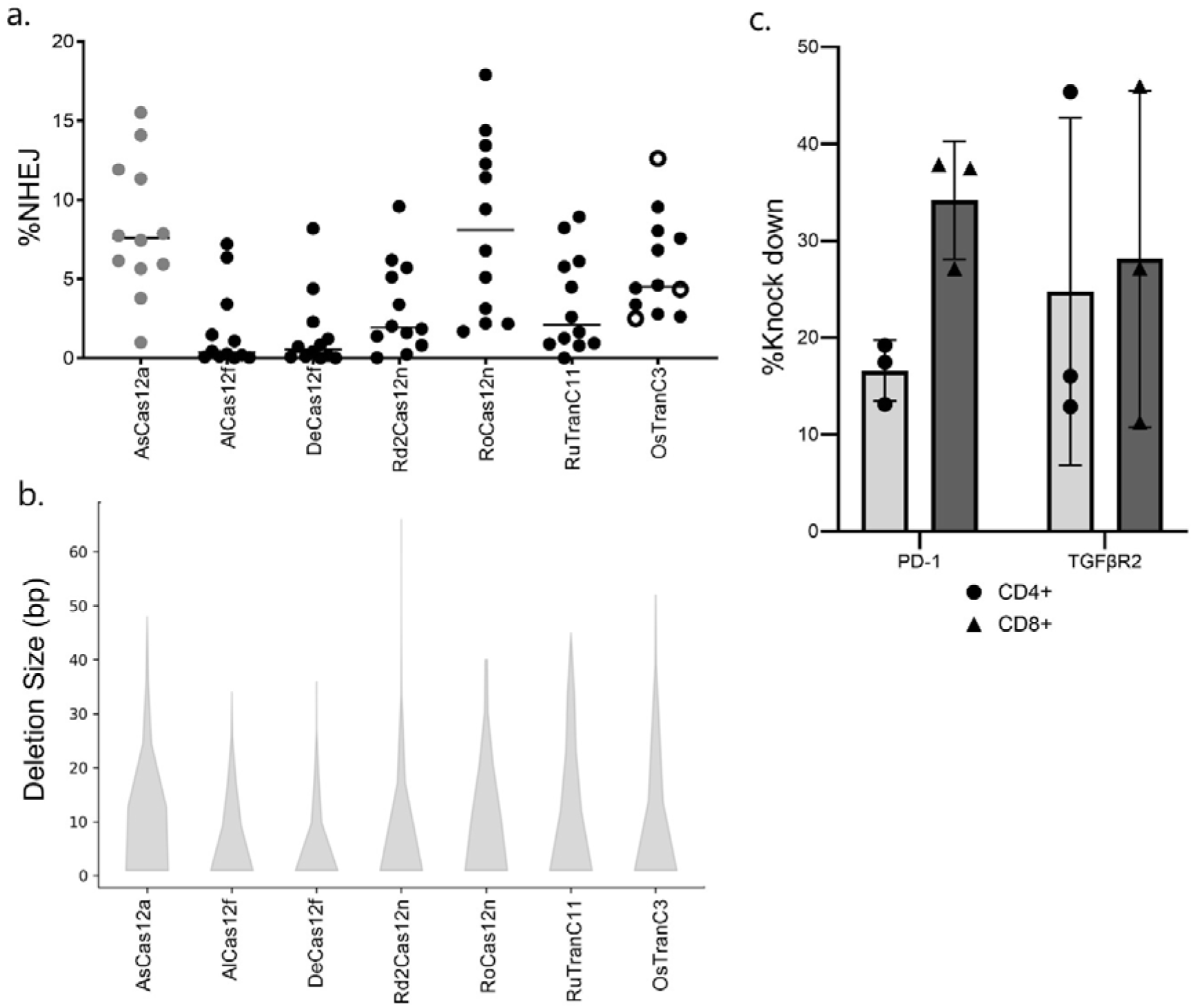
Eukaryotic performance of compact RNA guided nucleases. a. Non-homologous End Joining (NHEJ) activity cells of lead candidates and AsCas12a. Open circles indicate TTTC PAMs for OsTranC3. b. Size of deletions caused by lead candidates and AsCas12a. Data is from delivery in HEK293T cells at 12 targets, delivered in triplicate. All systems were tested on targets containing their respective identified PAM sequences. c. Functional knock of immunological regulators of T cell activity with OsTranC3. Each dot represents a separate nucleofection.

We observe that though AsCas12a typically outperforms compact systems, RoCas12n functions comparably to AsCas12a, while possessing a purine-rich PAM and being only 528 amino acids long. Another system, belonging to a novel subtype, OsTranC3, while performing slightly worse than AsCas12a, still possesses a notable level of activity for a compact system. The deletion profile of each of the leading candidate systems showed a difference in the cleavage patterns and median deletion for each type of RGN. The most common start position for deletions was 11-14 bp downstream of the PAM for AsCas12a, 23-25 bp downstream for the Cas12f systems, 13-16 bp downstream for two Cas12n systems, 16-20 bp downstream for OsTranC3, and 11-14 bp downstream for RuTranC11 (Supplemental Figure 7). The size of the resulting deletions was also smallest for Cas12f systems and Rd2Cas12n (median of 4 bp deleted), whereas RoCas12n (median=8), OsTranC3 (median=6), and RuTranC11 (median=8) had deletions of the same size as AsCas12a (median=7) (Figure 3b).

We then sought to further test one of the novel subtypes that was concurrently identified as a TranC and used our novel representative, OsTranC3, to target genes involved in immunological inhibitor signaling in T cells^12^. The PD-1/PD-L1 immune checkpoint is one of the most well-known T cell inhibitory pathways, where T cell activation results in upregulation of PD-1 and an intracellular signaling cascade that eventually leads to T cell suppression^21,23,24^. T cell effector functions are also suppressed by the regulatory cytokine transforming growth factor beta (TGF-β) which suppresses the immune response to solid tumors through TGFβ receptor 2 (TBGβR2)^20,22,25,26^. Accordingly, we measured the performance of mRNA delivery of the protein targeting PDCD1, and TGFβR2 in activated pan T cells. We directly measured the surface expression of PD-1 and TGFβR2 via flow cytometry and saw successful knock down of each protein (Figure 3c).

#### Specificity

To compare the performance and specificity of our leading candidates more directly, we utilized the recently published GenomePAM methodology^74^. This approach uses repetitive genomic sequences to enable direct comparison of enzymes with different PAMs at the same target sequence. Random flanking regions of the repeats are used for PAM targeting and characterization, and the degenerate nature of the repeats is used to measure specificity with output similar to that of GUIDE-seq^75^.

Slight differences between GenomePAM results and reported PAMs are known to occur, as it is a positive selection assay within mammalian cells, and the comparison to *in silico*, bacterial, and *in vitro* cleavage assays all occur under different conditions that may impact results^74^. We compared the compact editors to ISDRa2 TnpB with the optimized ωRNA-v2 sequence and to AsCas12a^76^. The PAMs identified via the eukaryotic assay generally agree with those identified via bacterial depletion (Figure 4a). We observed the canonical TTTV PAM first identified for AsCas12a through bacterial depletion studies^17^. We also see a TAM (ANTTGAT) that almost matches what was reported for ISDra2 TnpB (TTGAT), that was obtained through in vitro cleavage without the optimized ωRNA-v2 sequence^6,76^. The extra adenosine at position -7 does not appear as strongly for the PAM identified through mismatches, nor does the -2 adenosine (Supplemental File 1). OsTranC3 also demonstrates a slightly different PAM (TTTN vs TTC) than determined through bacterial depletion, however we observed activity on both TTTC and VTTC PAMs when tested on different targets, with no obvious difference in activity by PAM sequence (Figure 3a). The mismatch PAM shows a much weaker reliance on the initial thymine from the GenomePAM results than the next two, but still no preference for a final cytosine over other bases (Supplemental File 1). The GenomePAM workflow also allows for the direct comparison of system specificity, as the target between systems is held constant, thus removing its effect from consideration (Supplemental File 2). We compared the effect of increasing target specificity across the systems, by identifying reads that exactly matched for the specified target length (Figure 4b). As expected we see a higher specificity for AsCas12a compared to ISDra2 TnpB, with specificity decreasing with longer target sequences, matching the reported 16bp target length requirements for ISDra2 TnpB^6,77^. We also see a range of specificity for the rare and compact systems, with RoCas12n outperforming AsCas12a, and demonstrating the reported longer target sequence requirement for Cas12n relative to TnpB^27^.

**Figure 4.**
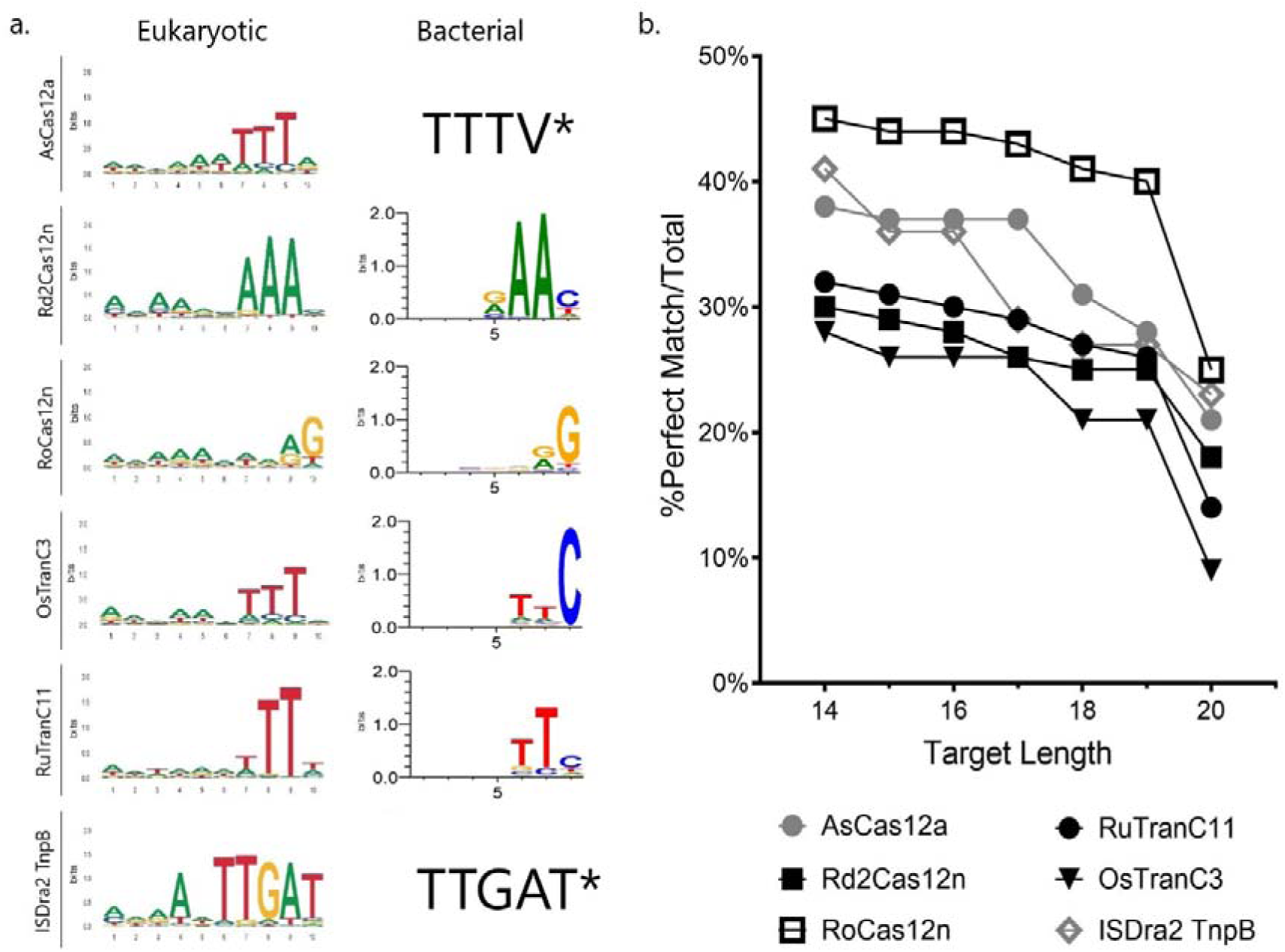
Eukaryotic PAM Recognition and Sequence Specificity. a. Recognition Sequences for Select Systems Determined through Eukaryotic Enrichment or Bacterial Depletion. *Bacterial PAM for AsCas12a taken from Zetsche et al, 2015^17^. Biochemical TAM for ISDra2 TnpB taken from Karvelis et al, 2021^6^. b. System specificity with increasing target sequence stringency of select systems.

RuTranC11 demonstrate a longer target specificity length than TnpB, similar to the longer target requirement Cas12n and Cas12f^78^. OsTranC3 however demonstrates a slightly longer target specificity length than TnpB, but it is shorter than the specificity lengths of the other compact systems.

## Discussion

The PAM of an RGN is an intrinsic property of the protein that determines what regions of the genome an enzyme can be programmed to target and provides the ultimate fidelity of the enzyme^13^. This has led to the search for more enzymes with unique PAM sequences to enable therapeutic editing of any SNP, while maintaining high on-target activity^60^. However, the two most abundant Class 2 (i.e., single-effector) RGNs in bacteria are ill-suited to form the basis of a therapeutic toolbox.

Cas12a is too large to be packaged into an AAV and does not possess diverse PAM sequences^4^. While Cas9 enzymes do present a variety of PAM sequences, they are generally large as well, typically longer than 1000 amino acids^69^. These systems have been co-opted for their DNA targeting ability to regulate transcription (CRISPRa and CRISPRi), target specific bases for modification (Base Editing), and insert specific sequences (Prime Editing or Programmable Gene Insertion)^1,3,60,79,80^. Fusing the effector domains for these other applications exacerbates the size problem by creating even larger constructs, posing a challenge for AAV-mediated therapeutic delivery. While split-AAVs have been developed for therapeutic applications, they have significant downsides, such as increased cost and immunogenicity^81^. Smaller CRISPR enzymes, such as Cas12f, allow for easier packaging into an AAV; however, the rarity of these systems has led to limited investigation into them, requiring faster and more efficient search strategies^68^. Highly abundant IscB and TnpB systems have been reported that are similar in size to other compact systems^5,6^. However, these systems have much more complex TAM requirements and lower target specificity requirements than CRISPR systems, limiting their applications^28,31,71,77,78.^

To more efficiently identify compact and rare systems that meet all the requirements for an ideal therapeutic platform, we conducted a search from a structure-focused perspective. Searching for nucleases in this way, we were able to yield hits that would have been missed by the usual approaches based on sequence similarity. Our search identified far more potential candidates in public databases than we were able to test, highlighting the richness and diversity of these enzymes, without the need to sequence deeper or from more exotic locations.

Protein structure predictions continue to improve in quality and decrease in computational requirements. As more public databases of folded proteins become available these provide a rich resource for future searches^82,83^. However, one of the main challenges we encountered was the difficulty in tracing the source genomic DNA for many hits from the ESMAtlas. This highlights the need for stronger data stewardship and governance, particularly with regards to cross-referencing data across multiple database types. This will become increasingly important as the amount of related heterogenous data collected increases exponentially. Ensuring best practices with data management allows the data to be further used and analyzed, enabling subsequent use cases that the original creators may not have foreseen.

While our search was limited to using the ancestral RuvC domain, future applications of this method could use alternative domains that are present in these nucleases, such as the HNH in Cas9, or the WED or REC domains, or be focused on RuvC folds of specific effector proteins. This search method could be further strengthened in several ways, such as examining the co-occurrence of signature domains in proteins, or employing more sensitive RGN-specific hidden Markov models derived from diverse multiple sequence alignments of structurally similar proteins. Finally, machine learning methods, such as the use of embeddings from protein language models that capture structural and functional information, could be used to further enhance the sensitivity of the search.

The search methodology we employed is scalable, making it compatible with the ever-growing amount of metagenomic data and allowing the identification of both rare and novel compact CRISPR subtypes. This allowed us to obtain enough of these systems to identify relevant tracrRNA characteristics applicable across several compact subtypes, enabling the rapid characterization of multiple enzymes in *E. coli.* Being able to separately express the gRNA and a codon-optimized version of the protein increased the success rate of PAM identification, making it comparable to that of the well-known Cas9 or Cas12a subtypes, which is critical to building a functional gene editing therapeutic tool kit^4^.

It is well known that many systems have some activity in eukaryotic systems but only a few have sufficient editing activity to be useful and may still require engineering to reach necessary activity levels^3,4,73^. Accelerating the process of identifying active systems consequently simplifies the search for useful systems. In this study, we identified six novel RGN subtypes that were concurrently identified as clades of the TranC family and subsequently deeply characterized multiple compact enzymes that demonstrate notable editing levels with only minimal gRNA optimization and demonstrated successful knockout of immunosuppressive barriers known to be active in solid tumors.

Our characterization of these novel compact editors also revealed unique cleavage patterns, specific to each subtype, confirming they are functionally distinct. These patterns show that the TranC systems preferentially cleave the target sequence at roughly the same position as AsCas12a and with a similar median deletion size.

Cas12n systems cleave consistently slightly further down the target sequence than Cas12a, as previously reported, while having notably different median deletion sizes^27^. Consistent with previous reports, Cas12f systems cleave outside the target sequence with a smaller median deletion size than AsCas12a^57^. Unlike TranC Clade 11, TnpB, and Cas12n, which have all been reported to function as monomers, Cas12f functions as an asymmetric homodimer, leading to its unusual cleavage location^12,70,71,78^. OsTranC3 predominantly cleaves at a location intermediate between Cas12f and the other reported RGNs of this study, so it remains to be seen whether it functions as a monomer or homodimer.

Our characterization also revealed differences in PAM calls between the typical negative in vitro or bacterial selection and positive eukaryotic enrichment^4,6,17,74^. We saw activity on PAMs identified via bacterial depletion that only partially overlapped with the PAM identified via positive eukaryotic enrichment with OsTranC3.

Additionally, previous reports from ourselves and others have also identified that AsCas12a can target non-canonical PAMs that were not identified through bacterial or eukaryotic PAM identification studies^4,17,84^. There have also been reports on disagreements of the PAM calls for CjCas9 through different methods^74^. These differences could be attributed to specific assay conditions, subjective calls of depletion, target selection, gRNA design, as well as differences in genome accessibility in eukaryotic cell lines. Additionally, the GenomePAM workflow does not have a truly random and even distribution of all possible PAM sequences, which could begin to affect the results, especially with long and complex recognition sequences, such as the TAM of ISDra2 TnpB^74^. All these variables stress the importance of proper and unbiased off target characterization of therapeutic targets to ensure patient safety.

Furthermore, we showed that it is primarily the Cas12a subtype that suffers from a lack of PAM complexity, and not all Type V or TranC effector proteins. The enzymes identified in this study and those used to compare recognition sequence diversity resulted from unbiased searches, so they should be representative of typical systems^4,5,12,29,30,69^. Novel Cas12 subtypes, arising from convergent evolution of disparate TnpB ancestral systems into true CRISPR systems, retain the TAM diversity seen among TnpB proteins but gain greater target sequence specificity and less complex PAMs. This discovery offers a key insight into the evolution of ancestral transposons into true CRISPR-Cas systems. For phage defense it is critical that the CRISPR-Cas protein not target the host genome, necessitating greater target specificity, but the system must be flexible enough in its PAM to identify multiple potential spacers that are targetable within the compact phage genome, requiring a less complex PAM. In contrast, in the “peel, paste, and copy" mechanism of TnpB, the TAM provides the recognition site for the transposon, and the target sequence directs TnpB to re-cleave the donor joint^6^. In this scenario, lower specificity of the target site could increase the ability to maintain the transposon at the original site after it has spread to a new location, while the complex TAM prevents collateral off-target cutting of the host genome^85^.

The evolution from MGEs to CRISPR-Cas systems also requires the creation and maintenance of the CRISPR array by other Cas accessory genes^14,18^. While our compact systems did not show associated accessory genes, this could be due to the fragmented nature of the metagenomic input data, limited contig length, or trans-encoded accessory genes located elsewhere in the genome. This fragmentary metagenomic data also highlights the strength of our structure-based search compared to other approaches which look for homology across a range of proteins in the CRISPR operon. With more fragmented genomes it is increasingly likely that a complete operon will not be contiguous and will thus be more difficult to identify through homology-based approaches.

This evolutionary switch from a purely selfish-gene-driven mobile genetic element (MGE) into a bacterial host defense system is still an area of active research^14,18,54^. It has been reported that Type V CRISPR systems first evolved from transposons through a functional RNA splitting of the transposon-derived right-end RNAs into crRNA and tracrRNA^12^. Here we report a TranC system that does not possess any expanded domains like other TranC or compact Cas12 systems, demonstrating that the splitting of the transposon-derived right-end RNAs (reRNA) and the ability to perform as an antiphage defense arose before the expansion of protein domains^12,54^. However, this system still has a compact PAM, like the typical PAM of other larger TranC/Cas systems, which is noticeably more compact than the TAMs of TnpBs.

This system (CauTranC9) was also identified in a phage genome, indicting that phages have acquired these systems for superinfection exclusion rapidly in their evolutionary history or that phages may serve as the original vector for the transition from a selfish-gene-driven MGE into a bacterial defense system^48^. The domain accretion from compact ancestral proteins over evolutionary time is common to all Class 2 CRISPR systems^40^. The precise role of these expanded domains in these systems remains to be confirmed, but it is plausible that the expanded REC and WED domains contribute to the improved specificity and longer target sequence requirements of these compact effector proteins compared to TnpB ancestral proteins, just as the expansion of the REC lobe in the evolution from IscB to Cas9 increased target DNA cleavage specificity^31^. We observed that OsTranC3, which does not have expanded REC and WED domains, had similar target length requirements as TnpB, but that those with expanded domains (Cas12n and TranC11) demonstrated longer specificity constraints. These results support the hypothesis that a compact TAM of a TnpB is a necessary evolutionary filter on the journey to intermediary TranC systems, where a relatively small PAM is required for effective antiphage defense to protect against a range of pathogens, which are also rapidly evolving to avoid defenses (Figure 5). The typical TnpB has a longer and more complex TAM, more suited to their role as MGEs. However, those TnpBs with a simple TAM, are able to be co-opted into an antiphage defense rather, creating an evolutionary preference for certain TnpBs to become Cas12 enzymes. In contrast, all Cas9 effectors are thought to derive from a single IscB ancestor but possess a wide diversity of PAM sequences^54^. This could reflect a higher mutability of the PAM-interacting domain of IscB or Cas9 systems than seen in large Cas12 systems, which is reflected by the high median Euclidean distance of IscB TAMs and the larger distribution of targetability scores compared to TnpBs. Additionally, as TnpBs are under negative selection pressure to limit their host toxicity, TranC/Cas12 offer a better alternative for the development of a gene editing toolbox, as they possess both the preferred compact PAM necessary for efficient genome targeting and potentially have been subject to positive selection for efficient editing^13,85^. Our search strategy provides an efficient method to identify these promising gene editing candidates, though which may offer an easier path for further engineering to realize their full potential.

**Figure 5.**
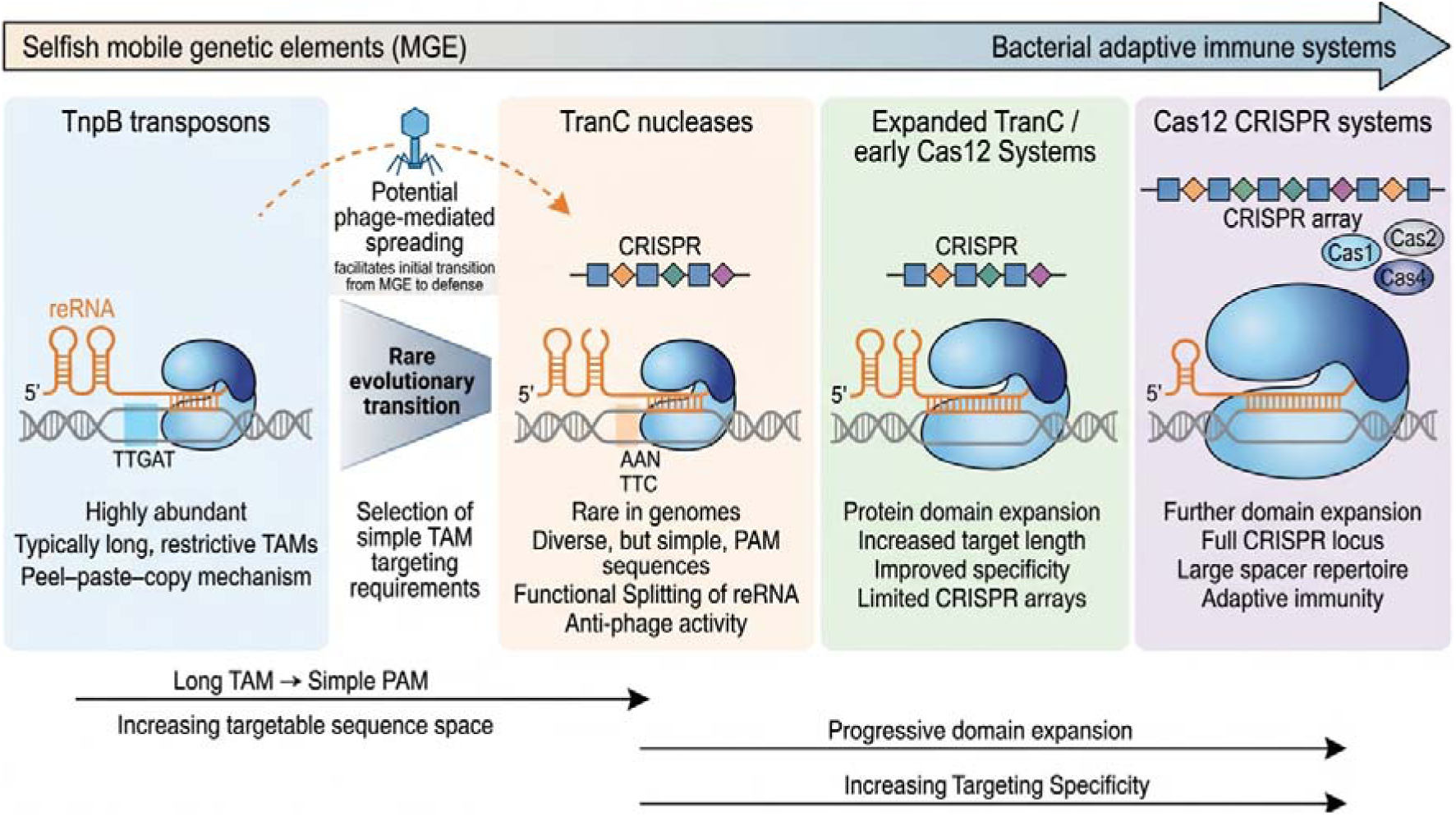
Evolution of TnpB transposons to Cas12 Bacterial Defense systems. A compact TAM provides an evolutionary filter for the development of Cas12 systems from ancestral TnpBs.

## Conclusions

Here we have shown that structural-based searches are capable of rapidly and efficiently identifying novel RNA guided nucleases with minimal sequence homology to the search query, which nonetheless maintain the same desired core activity. We show their use in a therapeutically relevant context to knock out immunological inhibitory signaling of activated T cells. We further show that CRISPR-associated RGNs are a better platform for developing a therapeutic toolbox, despite the much greater abundance of ancestral TnpBs and IscBs, due to their more flexible and compact PAMs and improved specificity. These systems elucidate the evolutionary path from ancestral TnpBs to intermediate TranC and finally full CRISRP-Cas systems, demonstrating an evolutionary filter for a simple PAM in TranC/Cas12 systems, in contrast to the complex TAM of TnpBs. These properties enable the identification of a system significantly smaller than Cas9 with a high genome targetability, while maintaining accuracy.

## Methods

### Sequence and structural alignment of proteins and RuvC domains

Multiple sequence alignments were generated using MAFFT with automatic algorithm selection^86^. PDB files for diverse RGNs were downloaded from the Protein Data Bank^55–59^. Structural alignments were performed using MUSTANG with rigid-body superposition enabled^87^. Pairwise root-mean-square deviation (RMSD) values in Angstroms were extracted from the alignments based on aligned Cα atoms and were converted to similarity using an exponential decay function. A permissive scale factor of 5 Å was used for the similarity function because of the low homology of the proteins.

### Prediction of structures for internal database

MinCED and PILER-CR were used to find CRISPR arrays in metagenomic bacterial, archaeal, and viral databases from gut and environmental origins^45–47,49,88,89^. Prodigal was used to identify potential coding sequences within 10 kb of a CRISPR array^90^.

Translated protein sequences were clustered at 90% sequence identity and filtered for length (>300 aa and <1150 aa). ESMFold was then used to obtain structural predictions of the filtered coding sequences^44^.

### Foldseek Query

Extracted RuvC domains were used in a Foldseek search against a Foldcomp-compressed full ESMAtlas database and a compiled database of ESMFold predicted protein structures described above, using the easy-search with “alignment-type 0” (local alignments)^50,91^. Hits in the database with an e-value < 1E-5 to at least one of the RuvC domains were considered for further analysis.

### Identification of CRISPR arrays near ESMAtlas Hits

The source DNA for hits from the ESMAtlas database was identified using the metadata associated with the database^44^. Briefly, the associated contig (MGYC) was identified from the predicted structure hit (MGYP), which was then cross-referenced with the EMBL-EBI MGNify database to obtain the source DNA^92^. Of the initial 6,337,508 MGYP hits, only 51% of the source contig references could be traced, and full DNA sequence was only obtained for 14% of the MGYP hits. CRISPR arrays located in the source DNA for ESMAtlas hits were identified as described above.

ESMAtlas hits that were near a CRISPR array were retained for further analysis.

### System Prioritization

Sequences greater than 1150bp in length or from the ESMAtlas database where the genomic DNA could not be identified were discarded^44^. CRISPR arrays in identified genomic DNA from ESMAtlas hits were identified using minCED, as was done with the internal database, and only those hits <10kb from a CRISPR array were retained. Sequences were deduplicated and divergence from known systems was determined by a search of the GenomeQuest database (GQLifeSciences, accessed November 2023). Sequences that were determined to be more closely related to TnpB than IscB and to contain an intact catalytic DED residue of a putative RuvC domain by MAFFT E-INS-I multiple sequence alignment (MSA) in Geneious Prime (Biomatters Ltd) were retained^5,29,86^. Known subtypes were determined by MAFFT E-INS-I MSA to known reference sequences (Supplemental Table 2). Where sampling location could be determined, sequences from extremophile environments were discarded.

CRISPRCasTyper was run on contigs >3 kb to eliminate sequences near an identifiable CRISPR-Cas operon and to identify any nearby accessory Cas proteins^33^. Systems that shared a high degree of sequence similarity but were not consistent in their position relative to the identified CRISPR array were discarded. A BLAST search of novel subtype sequences was performed on the NCBI database to identify further representatives, which were filtered in a similar fashion^93,94^. Only putative subtypes that were located upstream of their CRISPR array were selected for downstream characterization. A BLAST search of nearby ORFs was used to rule out the presence of TnpA from IS200/605 transposon family^5^. The most likely targets of the original crRNAs were identified using CRISPRTarget^95^. For systems not in NCBI, host taxonomy was identified using the MMSeqs2 taxonomy tool with the Genome Taxonomy Database^96,97^. Sequences without identified host taxonomy were designated as Environmental Genomic Sequence (EGS).

### Phylogeny Comparison

A multiple sequence alignment was performed on representative sequences (Supplementary Table 2) via using similar methods to phylogenetic trees generated for CasPEDIA^98^. Briefly, a multiple sequence alignment of the representative sequences was generated using MUSCLE^12,27,30,99^. Positions where more than 90% of the sequences contained a gap in the alignment were ignored. A tree was then constructed using iQ-Tree and, visualized using the Interactive Tree of Life v6, and rooted at the midpoint^100,101^.

### tracrRNA identification

Putative tracrRNAs for Cas12b and Cas12e subtypes were selected by identifying antirepeats between the end of the putative Cas protein and the start of the CRISPR array. A region upstream of the antirepeat of similar length to known examples was selected as a putative tracrRNA and joined to the crRNA with a GAAA tetraloop, and the predicted MFE was compared to validated examples in Geneious Prime (Biomatters Ltd)^64,65^. Putative tracrRNAs for Cas12n subtypes were selected by identifying the first antirepeat after the final catalytic RuvC residue of the putative Cas protein and the start of the CRISPR array with at least 6 of 10 bases pairing within the final 16 bases of the CRISPR repeat that was 100-200 bases in length, without including any bases upstream of the catalytic RuvC residue. The tracrRNA was joined to the crRNA with a GAAA tetraloop, and the predicted MFE was compared to validated examples in Geneious Prime (Biomatters Ltd). Previously identified Cas12n sequences, then called c2c9, were also searched for tracrRNA and characterized^8,27^. To identify the tracrRNA for Cas12f and novel compact systems, an antirepeat was identified between the start of the putative Cas protein and the start of the CRISPR array with at least 6 of 10 bases pairing within the final 16 bases of the CRISPR repeat, similar to previously reported methods^28^. 120-200 bases upstream of the potential antirepeat were then extracted and joined to the 5’ end of the region of the CRISPR repeat that bound to the antirepeat via a GAAA tetraloop. The MFE was then predicted in Geneious Prime (Biomatters Ltd) and trimmed to contain 4-5 stem loop sequences consistent with the gRNA diagrams for the Cas12n processed tracrRNA and the established Cas12f tracrRNAs. Representative structures were identified using aliFreeFold^102^. To improve activity of Cas12f and Cas12n, gRNAs were developed by truncating the stem-loops of the MFE diagrams in a similar fashions to those shown to improve activity in Cas12f and Cas12n^27,67,68^.

### Determination of PAM requirements for each RGN through Bacterial PAM Depletion

For systems where a tracrRNA could not be identified, as well as for the putative Cas12h system, the native CRISPR operon was synthesized under control of the T7 promoter in pET28 without any optimization. Three repeats of the CRISPR array were maintained, but the spacer sequence was replaced with a specific target sequence. Synthesis was performed by Genscript and Aster Biosciences. For systems with an identified putative tracrRNA, the identified Cas protein was codon-optimized for expression in *E. coli* via the Genscript codon optimization tool and synthesized with a NLS tag on each end under control of the T7 promoter in pET28^103^. Some Cas12n sequences were tested both with and without their unstructured carboxy-terminal tail, but no difference in PAM recognition was observed (data not shown)^27^. The identified targeting RNA with an appropriate targeting sequence was synthesized under the control of a separate T7 promoter on the same vector. All synthesis was carried out by Twist Biosciences and Aster Biosciences. PAM identification was performed as previously described, where a plasmid library with a target sequence flanked by a random 8mer library was delivered into cells containing an expression plasmid for the protein being tested^4^.

Active PAMs lead to plasmid cleavage and loss of antibiotic resistance and are depleted compared to controls. Amino acid sequences of codon-optimized active PAM systems, and their gRNA or crRNA are listed in Supplemental Table 1. Contig information of remaining active PAM systems without an identified tracrRNA or crRNA are also listed in Supplemental Table 1.

### Comparison of PAM diversity

The targetability score (TS) for each system was calculated by multiplying the relative frequency of each base or degenerate base in the recognition sequence and scaling by 1000. For example, TTTV would become 0.25*0.25*0.25*0.75*1000=11.72 TS.

Position weight matrices (PWMs) were calculated to represent the PAM preferences of each system. Only the probabilities of three nucleotides (arbitrarily selected) were included, as the fourth probability is overdetermined. Each PWM was padded to match the length of the longest PWM using “N” bases, i.e. equal probabilities of all nucleotides. Two-dimensional principal component analysis was run on an array created from the flattened PWMs, such that each row in the array corresponded to one PWM. A scatter plot was created from the two principal components.

### Dual Plasmid Lipofectamine Eukaryotic Activity Assay

Expression plasmids were synthesized for each selected enzyme with dual NLS tags after codon optimization for human expression (Genscript) into the pTwist CMV plasmid (Twist Biosciences)^103^. All Cas12n sequences were truncated to remove the dispensable unstructured carboxy-terminal tail^27^. The assay was then performed as previously reported, via dual plasmid delivery into HEK293T cells via lipofectamine transient transfection, followed by targeted amplification and next generation sequencing on a NextSeq1000 (Illumina) or an AVITI24 (Element Biosciences)^4^.

Results were analyzed with CRISPResso2^104^.

### Functional knockout of immunological inhibitory signaling

The codon-optimized sequence with dual NLS tags was synthesized as mRNA by Trilink using the M6 Cleancap and N1-methylpseudouridine base substitutions.

Guides were selected based on their proximity to successful guides used previously for functional knock out of PDCD1 and TGFβR2 and synthesized by Trilink^19^. Human T cells were isolated from purified PBMCs using the Pan T Cell Isolation Kit (Miltenyi Biotec) and activated using T cell TransAct (Miltenyi Biotec) according to the manufacturer’s instructions. 48 hours after activation, 1e6 cells were resuspended in P3 buffer (Lonza) along with 120 pM of total gRNA and 770 ng of effector mRNA before being nucleofected on the Lonza 4D Nucleofector using the EO-115 settings. Cells were then returned to recovery media with 1:1000 DNase supplemented and incubated for an additional 48 hours. The media was then refreshed and cells were incubated for a further 48 hours. Functional knockout of the receptors was confirmed by measuring protein levels via flow cytometry. Pan T cells were harvested and washed in FACS buffer (PBS + 0.05% BSA), then resuspended in FACS buffer containing cell surface antibodies and stained at 4C for 30 minutes. Cells were washed with FACS buffer and resuspended in FACS buffer containing DAPI at a final dilution of 1:50,000. Cells were then run on a Cytek Aurora Evo spectral flow cytometer and analyzed using FlowJo Software (BD Biosciences). Cell surface antibodies used for staining are as follows: CD3-BV785, CD4-PE Dazzle594, CD8-APCCy7, CD25-PE-Cy7, TGFβR2-PE, and CD279 (PD-1)-FITC (all from Biolegend).

### GenomePAM

The assay was carried out similar to the reported method^74^. Briefly, the plasmids for the effector protein and gRNA against target Rep-1RC plasmids were delivered into HEK293T cells along with annealed dsODN via lipofectamine. Four days later, gDNA was harvested and fragmented using NEBNext Ultra II FS DNA Library Prep Kit for Illumina (New England Biolabs) according to the manufacturer’s instructions, using a custom Y adapter. The sample was amplified with primers specific for the Y adapter (NEBNext Multiplex Oligos) and the dsODN template. The library was then reamplified using barcoded primers for sequencing on an AVITI24 (Element Biosciences) or NextSeq1000 (Illumina). The results were then analyzed using a modified version of the GenomePAM code that improved robustness and addressed some issues identified in the sequencing processing workflow^74^.

### Code availability

The modifications to the original GenomePAM pipeline have been submitted to the original GitHub^74^. The pipeline takes the output of paired NGS sequencing runs outlined in the methods as input.

### Data Availability Statement

The datasets analyzed during the current study are publicly available through the relevant published studies ^44–47,49^. The data supporting the findings of this study are available within the paper and its Supplementary Information or from the corresponding author upon reasonable request.

## Acknowledgments

The metagenomic data were produced by the U.S. Department of Energy Joint Genome Institute (https://ror.org/04xm1d337; operated under Contract No. DE-AC02-05CH11231) in collaboration with the user community. We thank Ashley Jermusyk^2^ and Lewis Scott^2^ for support, discussions, and help.

## Author contributions

ES, LR, and TB planned, implemented, and performed the structural search for novel Cas enzymes. MW and TB planned, executed, and performed the wet-lab screening experiments and gRNA improvements. TB, SC, and SH planned, implemented, and performed the functional T cell validation. LR performed the bioinformatics processing of the INDEL data. ES performed the phylogenetic comparisons and analyzed the Eukaryotic PAM data. All authors analyzed and interpreted the results of the wet-lab and computational experiments. MW, TB, ES, LR wrote the manuscript and SC and SH reviewed and edited the manuscript. TB conceived and designed the study. All authors have reviewed and approved the final version of the manuscript.

## Supplementary Data

Supplementary_Info.xlsx: Supplementary tables containing relevant data for this study gRNA

Supplementary Table 1 Novel RGN sequences, taxonomy, PAMs, and

Supplementary Table 2 RGN sequences used for Phylogenetic Tree

Supplementary Table 3 PAMs and TAMs used for comparison

Supplementary Table 4 aliFoldResults

Supplementary Table 5 CRISPRTarget Results for CauTranC9

Supplementary Table 6 NGS Primers for eukaryotic validation

Supplementary Table 7 gRNA used to test systems in Eukaryotes

Supplementary Table 8 Sequences used to test T cell editing

Supplemental File 1. GenomePAM Cumulative alignments’ read counts for each enzyme

Supplemental File 2. GenomePAM identified targets for each enzyme

---

**Supplemental Figure 1.**
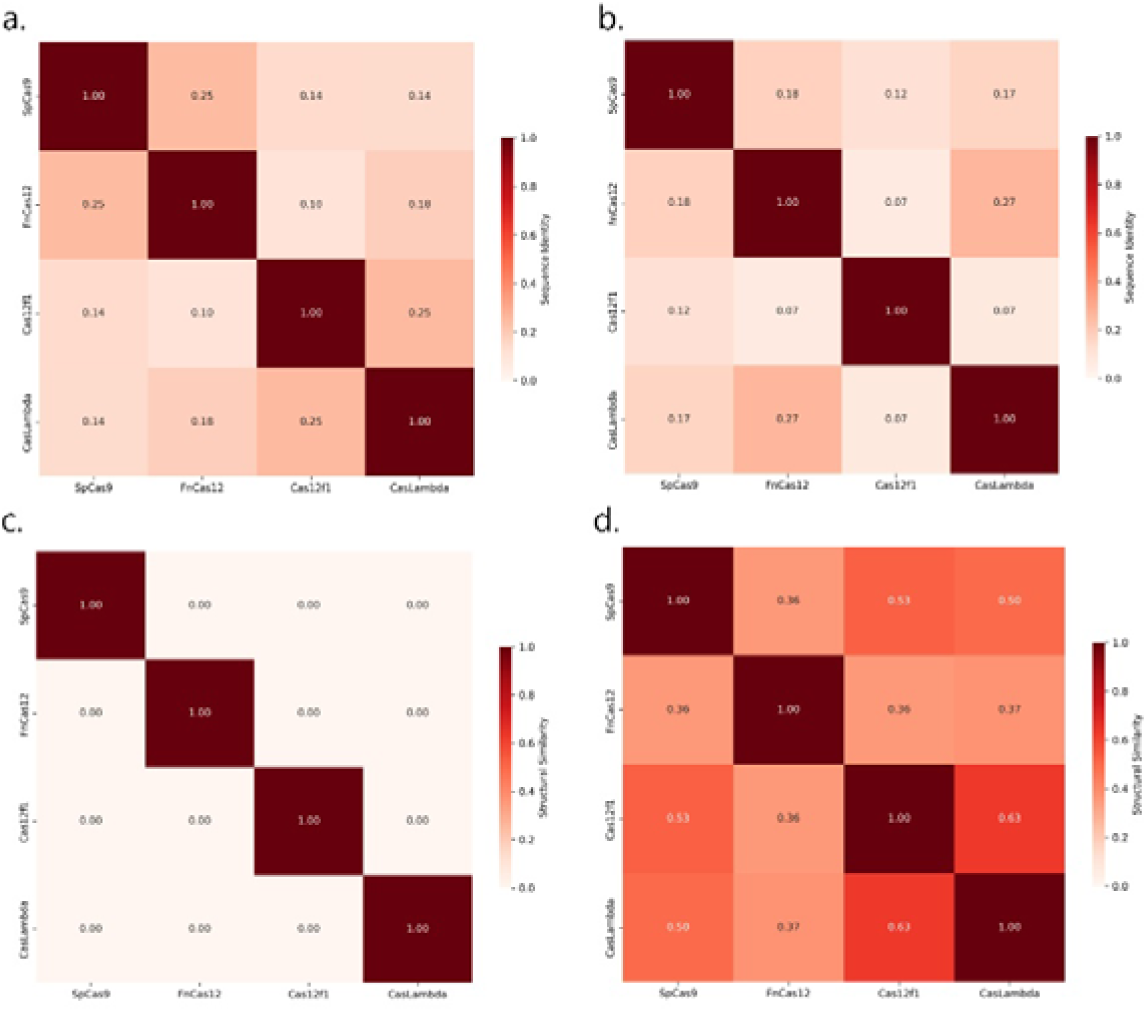
Sequence and structural comparison of search proteins. a. sequence identity of full length RGNs. b. Sequence identity of RuvC domain of search RGNs. c. Structural similarity of search RGNs. d. Structural similarity of RuvC domain of search RGNs

**Supplemental Figure 2.**
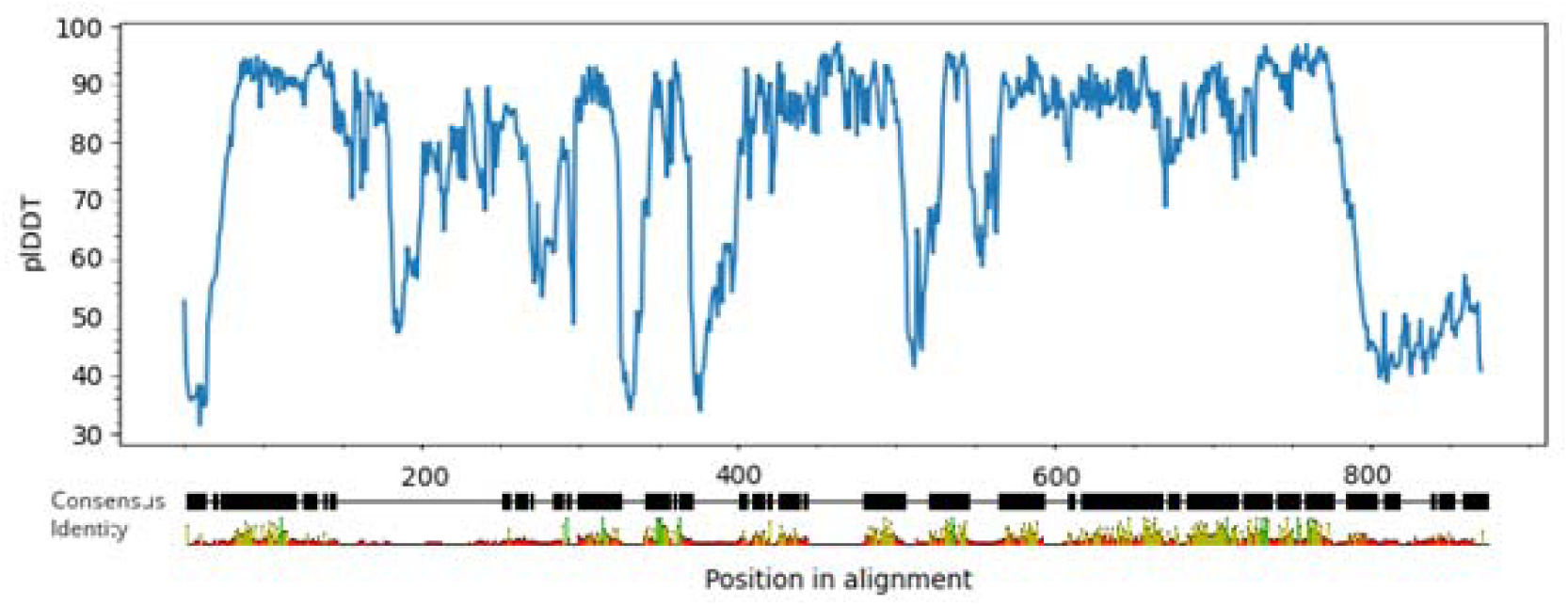
Consensus pLDDT values by position of aligned Cas12n sequences.

**Supplemental Figure 3.**
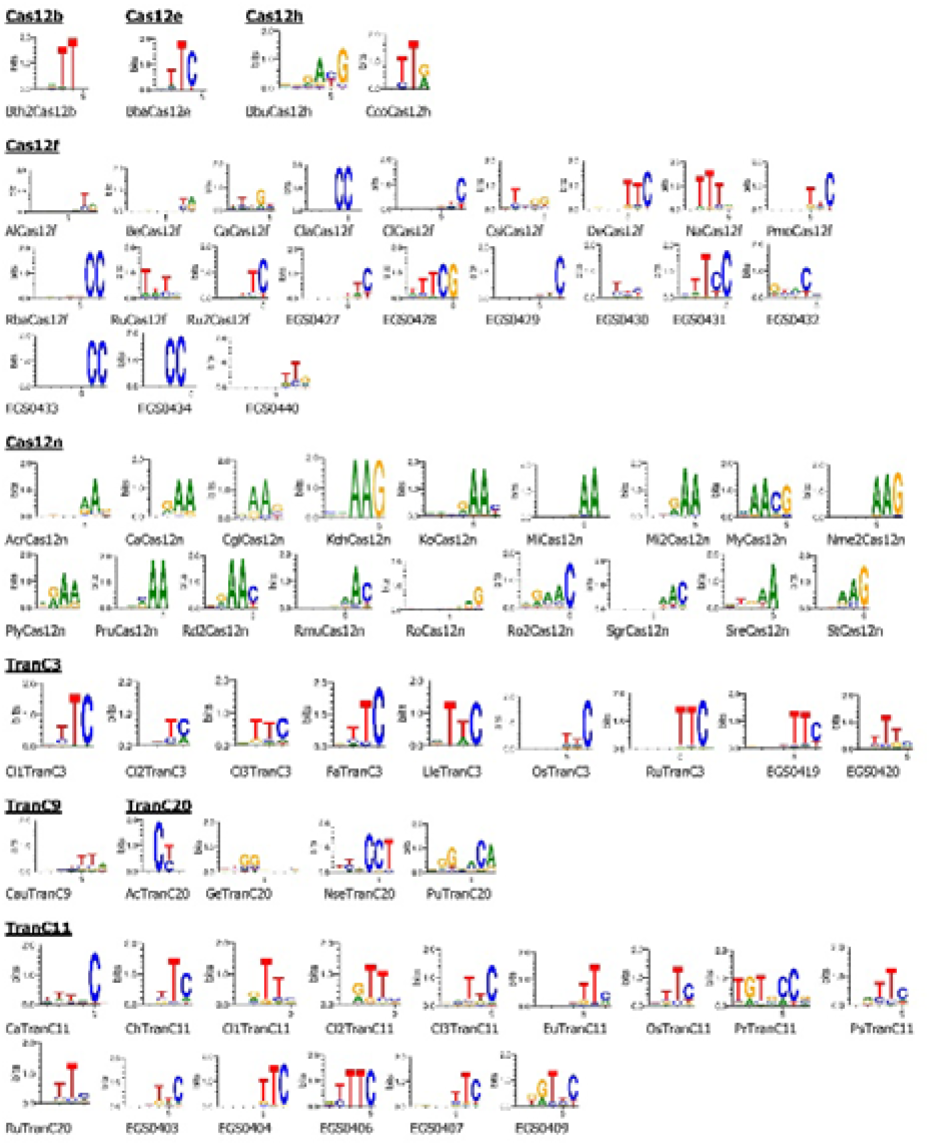
Identified PAMs of active rare and novel compact systems.

**Supplemental Figure 4.**
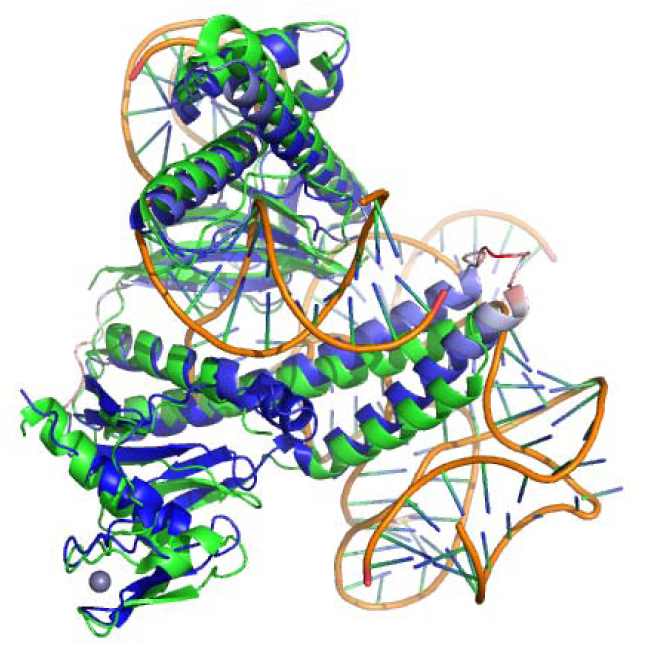
Predicted structure of CauTranC9 aligned to the structure of TnpB (green). pLDDT values range from 70 (red) to 100 (blue).

**Supplemental Figure 5.**
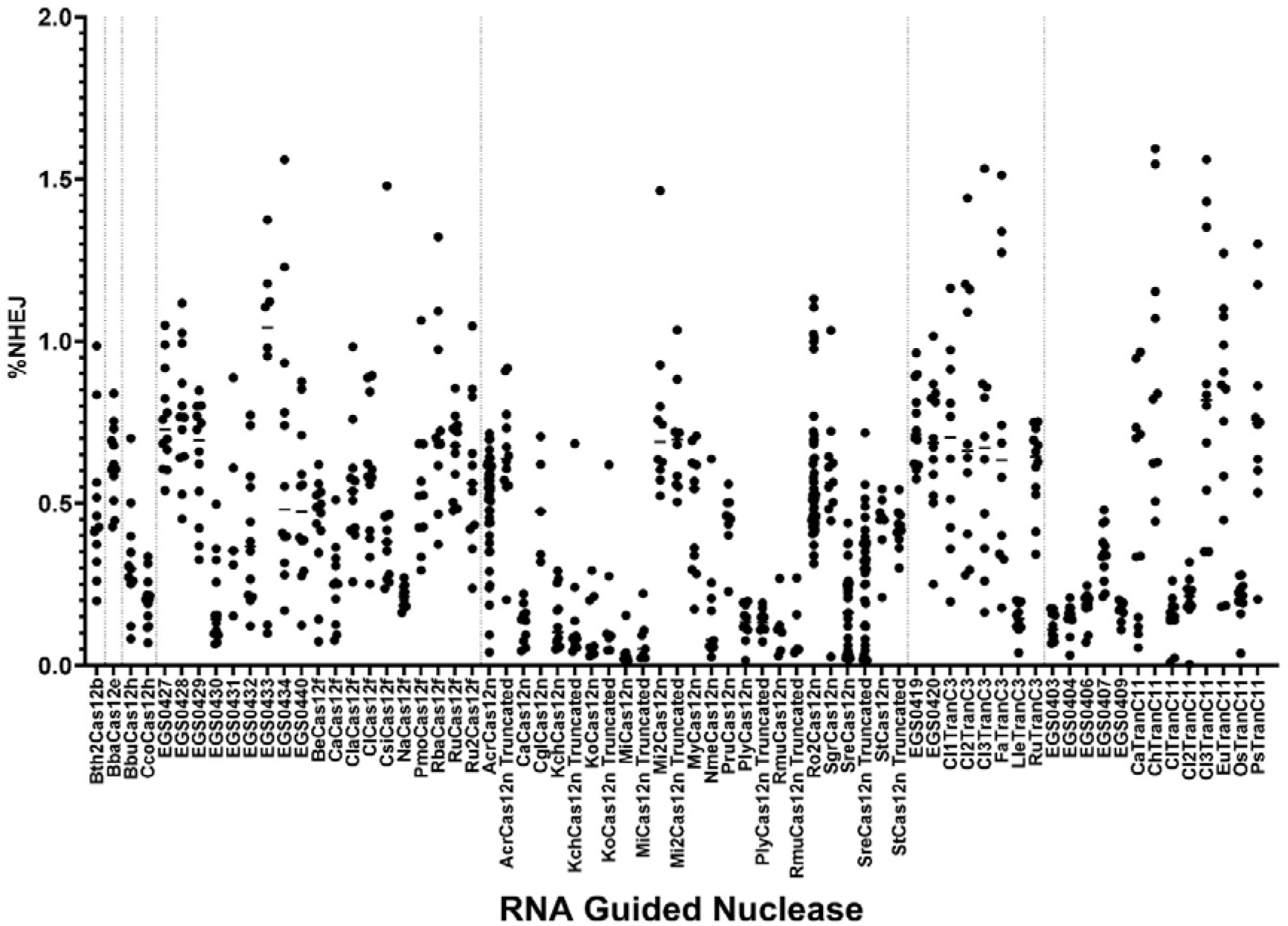
Eukaryotic editing of other nucleases advanced for testing from bacterial depletion. Each dot represents an independent target delivered 1-3 times. Dashed lines separate different protein families

**Supplemental Figure 6.**
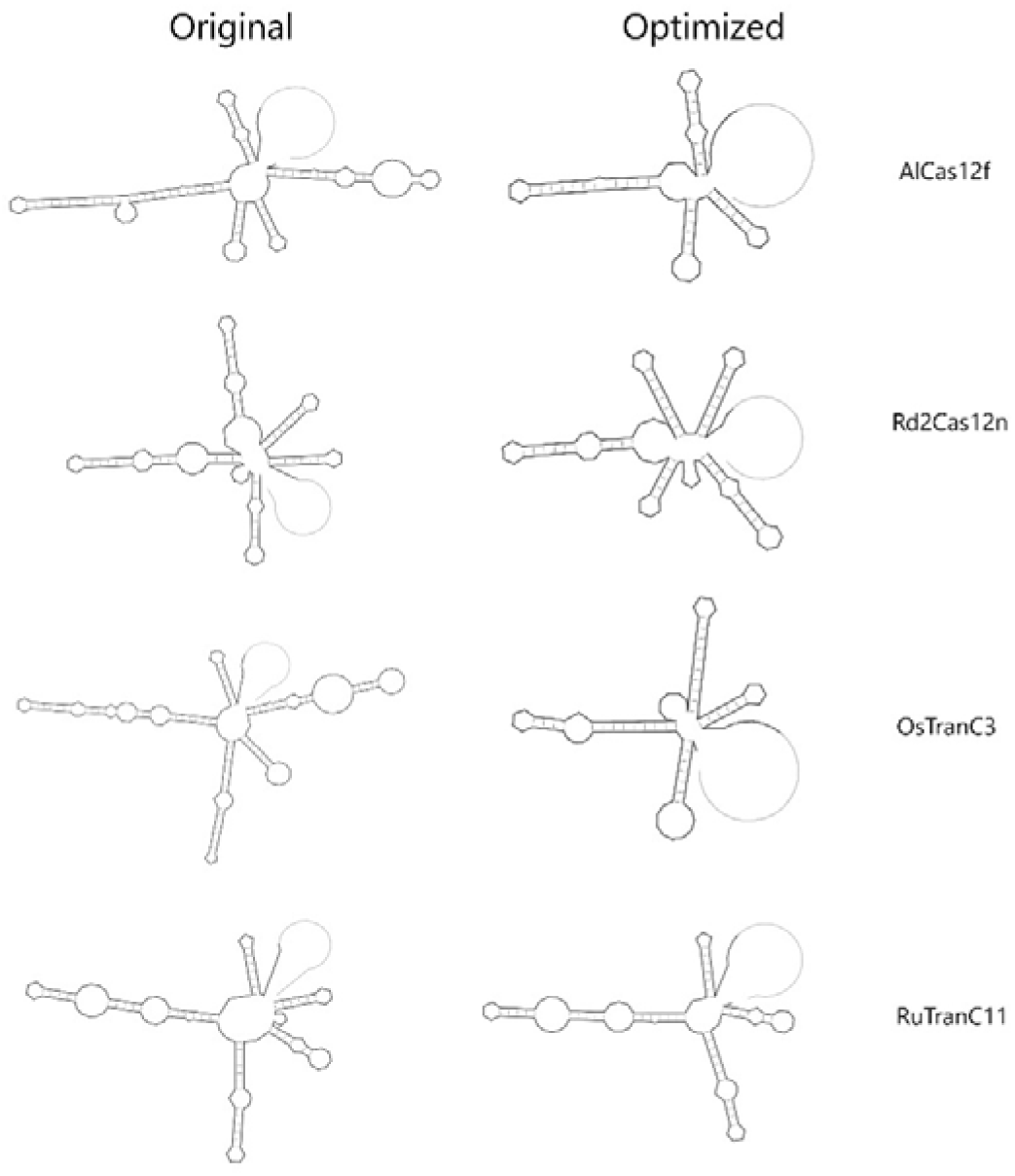
Improved gRNA designs of compact systems. Grey lines indicate reprogrammable targeting region.

**Supplemental Figure 7.**
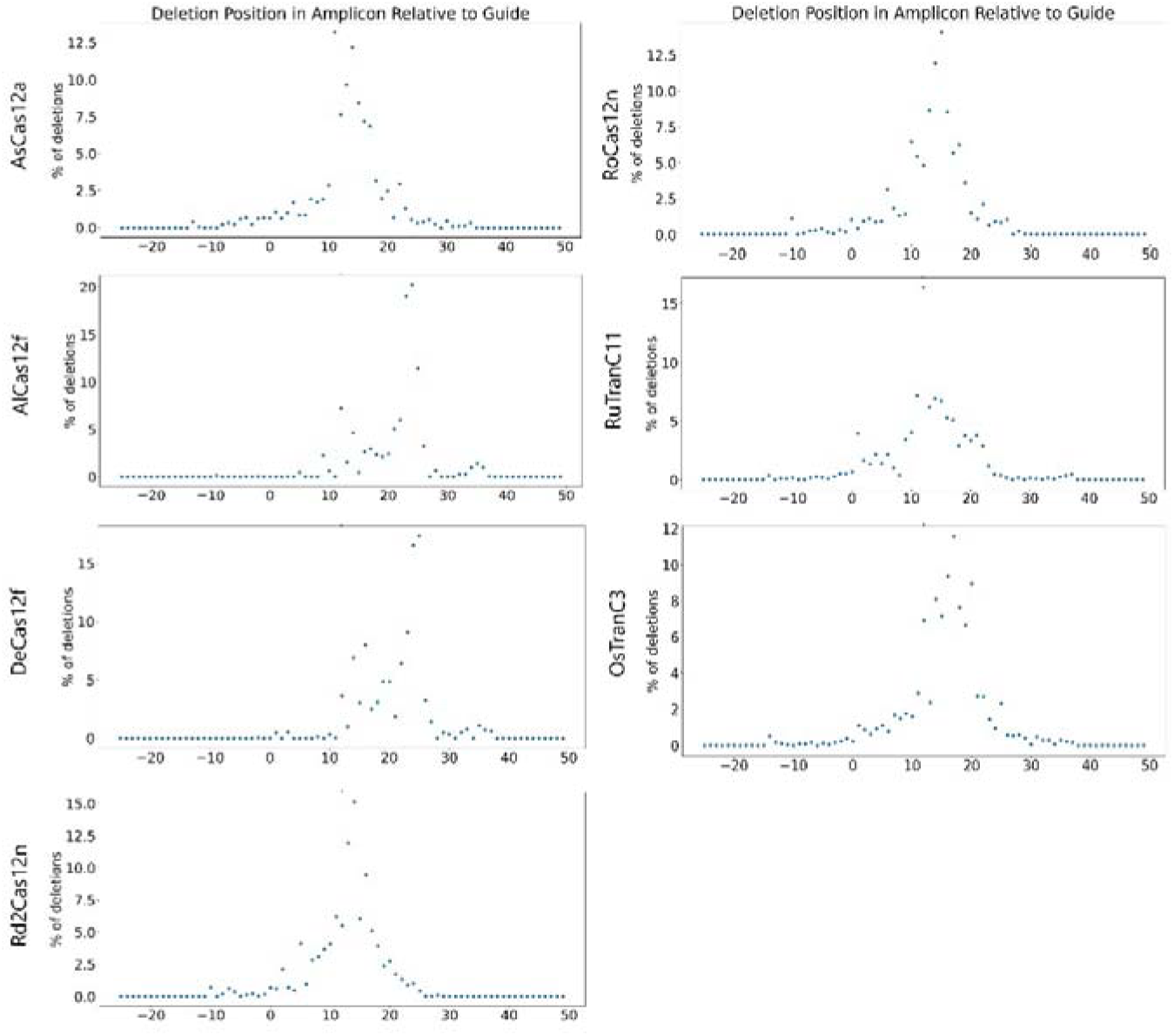
Start position of deletions in amplicon relative to guide.

